# Targeting primary and metastatic ovarian cancer with a peptide derived from the human NAF-1/CISD2 protein

**DOI:** 10.1101/2024.12.26.630413

**Authors:** Ehud Neumann, Yang Sung Sohn, Ola Karmi, Merav Darash Yahana, Alfredo E. Cardenas, Eli Pikarsky, Ron Elber, Assaf Friedler, Ron Mittler, Rachel Nechushtai

**Author notes:** These authors contributed equally: Ehud Neumann, Yang Sung Sohn.

## Abstract

Ovarian cancer is the most fatal cancer of female reproductive organs. Ovarian cancer is typically diagnosed at a late stage, after metastasis occurred, leading to a 5-years relative survival rate of only ∼5%. Here, we demonstrate the anti-ovarian cancer properties of a peptide derived from the human protein CISD2/NAF-1 (3D-NAF-1^44-67-6K^). This peptide selectively permeates the plasma membrane of ovarian cancer SKOV-3 cells without affecting healthy cells. 3D-NAF-1^44-67-6K^ targets and destroys the cancer cells’ mitochondria which leads to cancer cell death. *In vivo* studies of mice carrying xenograft tumours of SKOV-3 showed that the peptide significantly decreased the overall size and growth rate of both primary and metastatic ovarian cancer tumours. We further show that 3D-NAF-1^44-67-6K^ has a broad-spectrum anticancer activity targeting leukaemia, brain, and pancreas cancer cells. Our study suggests that 3D-NAF-1^44-67-6K^ could be used, alone or in drug combinations, to treat ovarian cancer and improve patient survival.

## Introduction

Ovarian cancer (OC) is the most fatal among female reproductive organs cancers^1, 2^. Epithelial OC is the most common type of OC and accounts for 95% of all OC cases^3^. Epithelial OC can be divided into five primary histologic subtypes: High and low grade serous carcinomas (HGSC and LGSC, respectively), endometrioid, mucinous, and clear cell^4^. Despite its name, OC can originate from non-ovarian tissues^5, 6, 7^. HGSC tumours usually originate at the fallopian tube and are found in the ovary after implantation^8, 9^. Non-serous (NS) tumours generally originate at the ovary and can migrate faster than the HGSC tumours^9^. A common feature of all OC is their localization to the ovary and/or related pelvic organs^10^.

Due to diagnostic difficulties, 80% of OC patients are diagnosed at stage IV-V when metastatic tumours already spread^11^. At this stage, the 5-year relative survival rate is only 5%^11^. Currently, the most common treatments of OC are surgery^12^ and chemotherapy^13^. Attempts to develop additional therapeutic approaches did not yet show promising results in most OC patients^13^. Even if OC is diagnosed early and treated, 80% of the patients experience relapse within 18 months^14^.

Mitochondria play an important role in cells mediating processes such as energy production, iron-sulfur cluster (Fe-S) biogenesis, apoptosis etc^15^. Mitochondria are also essential for cancer development, migration, and metastasis^16, 17, 18, 19, 20, 21^. Mitochondrial morphology is highly dynamic and can change via two main processes: fusion (elongation) and fission (fragmentation)^15, 16^. In cancer cells, the different morphologies of the mitochondria were associated with tumour behaviour. Mitochondrial fusion appears to support primary tumour growth and decreased metastasis^19^, whereas mitochondrial fission has been associated with increased metastasis^21, 22, 23^.

Due to the important role of mitochondria in cancer progression, mitochondria became one of the primary targets for anti-cancer drug design in different cancers^24, 25^. Conventional chemotherapy drugs that target mitochondria suffer however from harsh side-effects and/or the development of drug resistance^26, 27, 28, 29, 30, 31, 32^. Although derivatives of some of these drugs have been modified with relatively small compounds such as cationic molecules^33, 34^, microcapsules^33, 34^, and/or peptides^26, 27, 28, 29, 30, 31, 32^, to increase precision, newer drugs/drug combinations are urgently needed to treat OC.

The cancer cell plasma membrane (PM) differs in composition^35, 36, 37^, potential^38, 39^ and extracellular environment^40, 41^, from that of healthy cells. These unique properties of cancer cells make them an attractive target for drugs or peptides that selectively target them. Recently, we reported that a peptide derived from the transmembrane and disordered domains of the mitochondrial/ER protein CISD2/NAF-1 (3D-NAF-1^44–67^) can selectively target breast cancer cells without harming healthy cells^33^. Upon specifically permeating the breast cancer cell PM, the peptide induces several cell death pathways in cancer cells, including apoptosis, ferroptosis and necrosis^33^.

Here, we show that an improved version of our original peptide, *i.e.,* 3D-NAF-1^44–67–6K^, is a cancer-targeting peptide which selectively permeates the PM of SKOV-3 and OVCAR-3 OC cells and targets their mitochondria. 3D-NAF-1^44–67-6K^ causses mitochondrial fragmentation and decreases the SKOV-3 and OVCAR-3 cell viability. *In-vivo*, in a human xenograft mice OC model, 3D-NAF-1^44-67-6K^ decreases primary and metastatic tumour size and growth rates, without affecting healthy mice cells. These results suggest that 3D-NAF-1^44-67-6K^ could be used, alone or in drug combinations, to treat OC and improve patient survival. In addition, we show that 3D-NAF-1^44-67-6K^, can function as a broad-spectrum anti-cancer peptide that targets leukaemia, brain, and pancreas cancer cells. This finding highlights 3D-NAF-1^44-67-6K^ as a potential treatment for additional unmet cancer types that urgently need new drugs/therapies.

## Results

### 3D-NAF-1^44-67-6K^ displays enhanced cytotoxicity towards cancer cells

Addition of charged amino acids to a protein/peptide is known to increase its solubility in aqueous solutions^42^. To increase 3D-NAF-01^44–67^ solubility and uptake into cancer cells, six lysine residues were added to its c-terminus. The modified peptide was tested for its cytotoxic effect *vis-a-vis* the 3D-NAF-1^44–67^ peptide using control and cancerous epithelial breast cells (Supplementary Fig. 1). Both peptides decreased the viability of cancer cells after 1 and 3 days, compared to control non-cancerous cells that were not affected. However, compared to the 3D-NAF-1^44-67^ peptides, the improved 3D-NAF-1^44-67-6K^ peptide showed a significantly higher cytotoxicity towards cancer cells. To test whether the addition of the positive charge of the six lysine residues was responsible for the higher toxicity of the modified peptide, we replaced the positively charged 6 lysine tail with a positively charged 6 arginine tail (3D-NAF-1^44-67-6R^) and tested the cytotoxicity of the Arg-peptide (Supplementary Fig. 2). Compared to the 3D-NAF-1^44-67-6K^ peptide, the 3D-NAF-1^44-67-6R^ peptide had a lower killing activity (Supplementary Fig. 2). These results suggest that the addition of a positive charge contributed to the increased cytotoxicity of the peptide, however additional chemical or structural properties (that differentiate between Arg and Lys) might also contributes to the 3D-NAF-1^44-67-6K^ increased toxicity. Computational simulations, described in Supplementary Table 1 and Supplementary Fig. 3, suggested that, compared to the 3D-NAF-1^44–67^ peptide, the 3D-NAF-1^44-67-6K^ peptide interacted more with the cancer cells membrane during the early permeation events, leading to superior permeation properties into the breast cancer cells PM. This provides a possible explanation to the improved cytotoxicity of the modified peptide.

### Cytotoxicity of 3D-NAF-1^44-67-6K^ towards ovarian cancer cells

To study the toxicity of the 3D-NAF-1^44-67-6K^ peptide towards OC cells, we treated two different OC cell lines, SKOV-3 and OVCAR-3, with different concentrations of the 3D-NAF-1^44-67-6K^ peptide and measured their viability (Fig. 1 a-b).

**Fig. 1.**
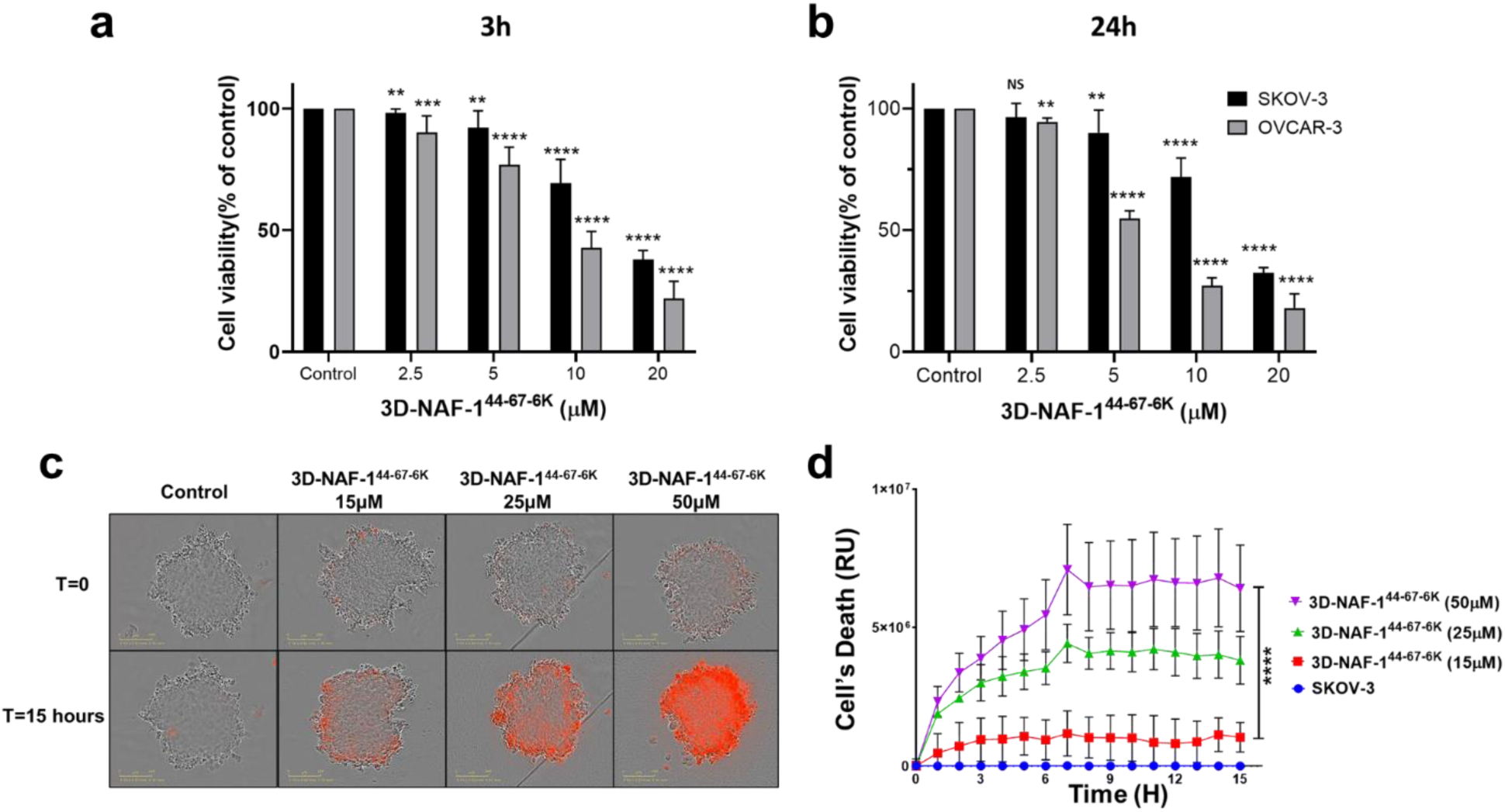
Toxicity of 3D-NAF-1^44-67-6K^ towards ovarian cancer cells. **(a-b)** Cell survival (precent) of SKOV-3 and OVCAR-3. Cell’s survival was measured at T=3h **(a)** and T=24h **(b)**. **(c)** Representative images of SKOV-3 spheroid 3D-cultures in the presence or absence of the NAF-1^44-67-6K^ peptide at the start of the measurement and after 15 hours. IncuCyte® Red cytotoxicity reagent was used to measure cell death (red signal). **(d)** The effect of 3D-NAF-1^44-67-6K^ on SKOV-3 cell death in spheroid cell culture over time. *\*\*P<0.01, *** P<0.001, **** P<0.0001* by t-test.

At the lowest concentration of 2.5μM, the peptide did not show a significant effect on the viability of both OC cell lines even after 24 hours. At the slightly higher concentration of 5μM, 3D-NAF-1^44-67-6K^ showed almost no cytotoxic effect towards SKOV-3 cells, even after 24 hours, while displaying significant toxicity towards OVCAR-3 cells (25% and 45% cell’s death after 3 and 24 hours, respectively). At higher concentrations of 10μM, after 3 hours the peptide decreased SKOV-3 cell viability by 30%, while OVCAR-3 cell viability decreased by 60% (Fig. 1a), resulting in almost double the amount of cell death. After 24 hours, the viability of SKOV-3 cells did not decrease further, while OVCAR-3 viability dropped to around 75% (Fig. 1b). At the highest concentration of 20μM, the 3D-NAF-1^44-67-6K^ showed high toxicity toward both OC cell lines. Higher concentrations of the peptide therefore led to a higher decrease in both OC cell lines viability, indicating does-dependent effect with an IC50 of 11μM and 15μM for OVCAR-3 and SKOV-3, respectively (Supplementary Fig. 4). These results indicate that 3D-NAF-1^44-67-6K^ is cytotoxic towards different OC cell lines. SKOV-3 cells were showed to be more invasive^43^, migrate at higher rates^9^ and are more resistant than OVCAR-3 cells^44^. We therefore decided to focus on the effects of the 3D-NAF-1^44-67-6K^ peptide on the more aggressive SKOV-3 cell line.

To farther investigate 3D-NAF-1^44-67-6K^ toxicity towards cancer cells, SKOV-3 3D-spheroids were grown and incubated with different concentration of the peptide (Fig. 1b-c). At a concentration of 15μM peptide, a relatively small amount of cell death was observed. Increasing the peptide concentrations to 25μM and 50μM resulted in increased cell death of about 3.7-and 6.2-fold, respectively.

### 3D-NAF-1^44-67-6K^ induces fragmentation of ovarian cancer cells’ mitochondria

To study the entry of the peptide into SKOV-3 cancer cells, and its effects on mitochondria, we labelled the peptide with fluorescein (Fl-3D-NAF-1^44-67-6K^), and the mitochondria with Rhodamine 800 (which is sensitive to membrane protentional), and imaged cells using confocal microscopy. Upon addition of the peptide to SKOV-3 cells, Fl-3D-NAF-1^44-67-6K^ fluorescence was primarily detected near SKOV-3 cells’ PM (Fig. 2a-b). At 4 and 6 hours of incubation, high Fl-3D-NAF-1^44-67-6K^ fluorescence was also detected inside SKOV-3 cancer cells (Fig. 2a-b). These findings indicated that Fl-3D-NAF-1^44-67-6K^ permeates SKOV-3 cancer cells. Following the permeation of the peptide into the cancer cells, the staining of mitochondria with Rhodamine 800 in treated cells decreased indicating that the peptide negatively impacted mitochondrial function (Fig. 2c). In contrast, the fluoresceine dye applied by itself (Fluorescein) did not have any effect on cancer cells and/or the mitochondria (Fig. 2b-c and Supplementary Fig. 5). This effect was also observed using another fluorescence probe, Cy5 conjugated to 3D-NAF-1^44-67-6K^, whereas no effect on mitochondria was observed with Cy5 alone, as well as in controls without treatment (Supplementary Fig. 6).

**Fig. 2.**
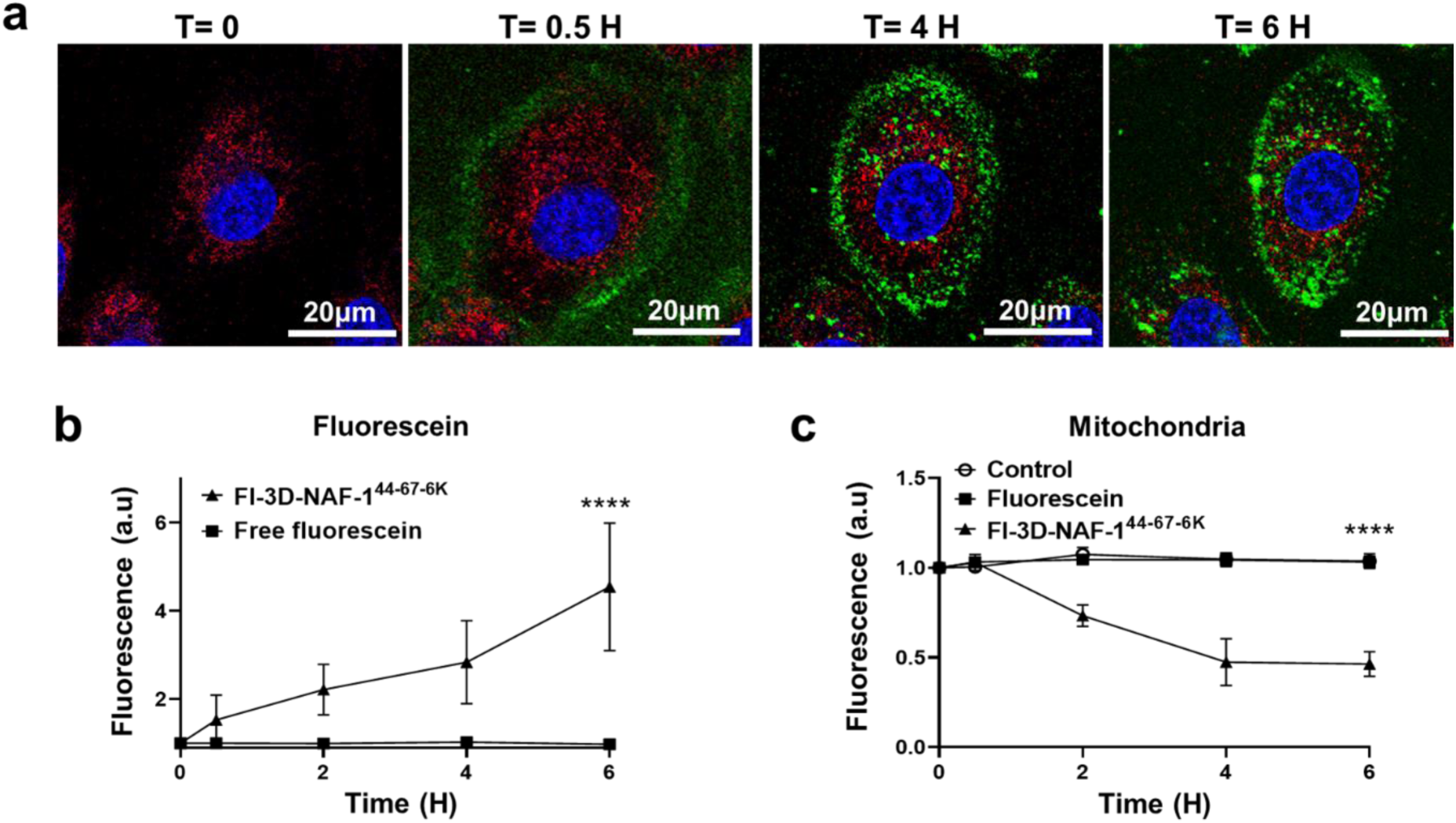
Permeation of FL-3D-NAF-1^44-67-6K^ into ovarian cancer cells and targeting of mitochondria. **(a)** Representative confocal fluorescence images of SKOV-3 ovarian cancer cells at different time points following treatment with Fl-3D-NAF-1^44-67-6K^ peptide. **(b-c)** Quantitative analysis of Fl-3D-NAF-1^44-67-6K^ and free Fluorescein permeation into the SKOV-3 cells **(b)** and mitochondrial damage, as accessed by mitochondria membrane potential loss imaged with Rhodamine 800 **(c)**. Data are shown as mean ± SD of 15 cells per field (10 fields in total) calculated for each time-point obtained from 3 independent experiments. *\*\*\*P < 0.001* by t-test.

The effect of the 3D-NAF-1^44-67-6K^ peptide on the mitochondria of SKOV-3 cells was further investigated by measuring the length of mitochondria in SKOV-3 cells at different concentrations of the peptide (Fig. 3; Supplementary Fig. 7).

The peptide therefore showed a dose-dependent effect on cancer cell mitochondria that included a decrease in mitochondrial length and membrane potential (Figs. 2 and 3). This result indicates that the 3D-NAF-1^44-67-6K^ peptide targets the mitochondria of SKOV-3 cells, causing mitochondrial fragmentation and loss of function.

### The 3D-NAF-1^44-67-6K^ peptide significantly decreases the migration of OC cells

Migration of cancer cells is associated with increased metastasis^21, 22, 23^. We therefore, measured SKOV-3 cancer cell migration in the presence or absence of the peptide for 96 hours (Fig. 4 and Supplementary Movie 1-3). SKOV-3 cells were planted on the edges of the well and allowed to migrate freely in the presence or absence of the peptide. The percentage of the areas taken by the cells was then measured at each time point. At a low concentration of 5μM, 3D-NAF-1^44-67-6K^ showed a minimal effect on SKOV-3 cells migration, compared to control. At 10μM, 3D-NAF-1^44-67-6K^ decreased SKOV-3 migration by about 30% compared to control. At the highest concentration of 20μM, 3D-NAF-1^44-67-6K^ showed the highest decrease in SKOV-3 migration of about 73%, indicating a dose dependent effect of the peptide on SKOV-3 cells migration.

**Fig. 3.**
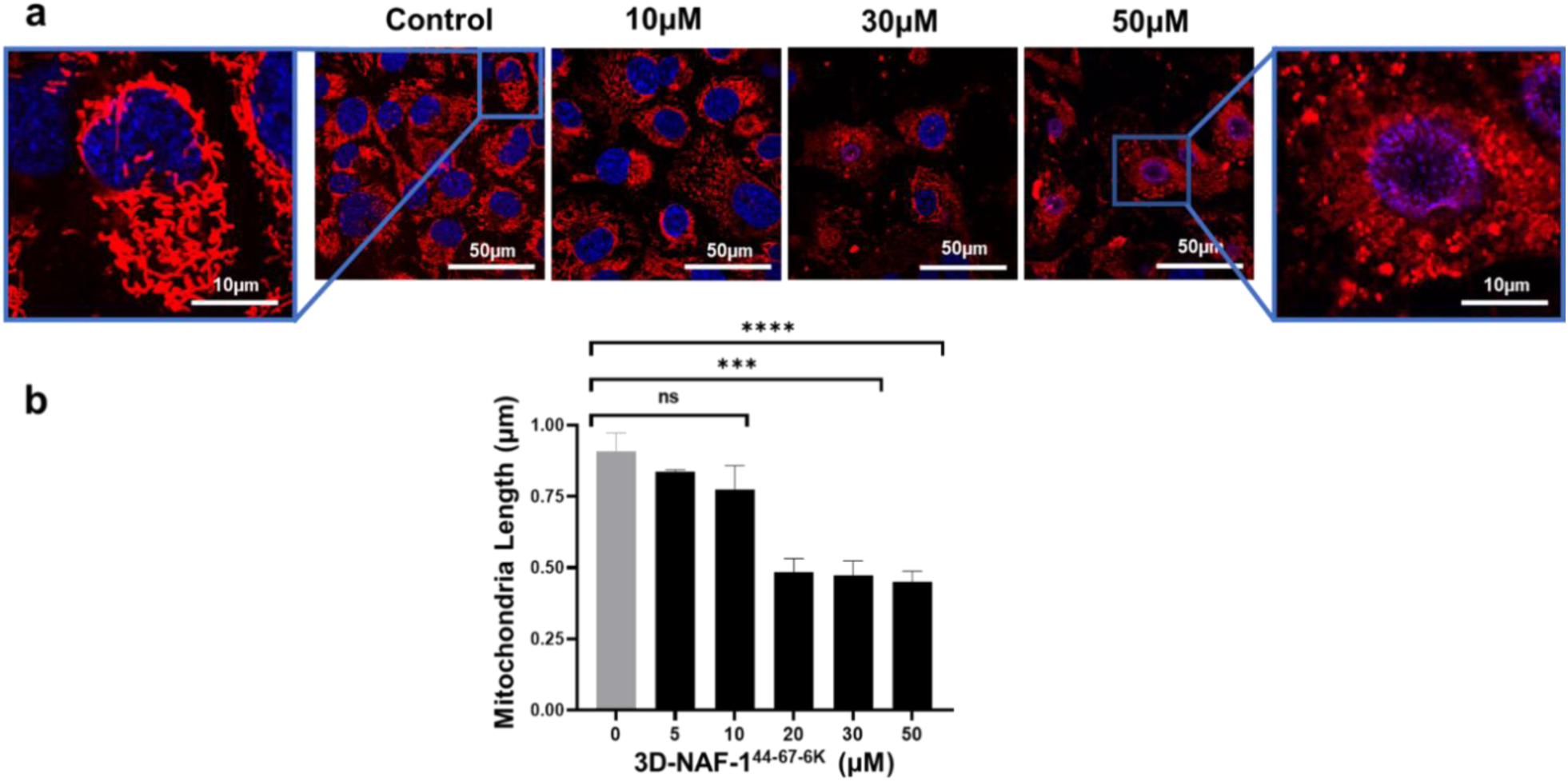
The 3D-NAF-1^44-67-6K^ peptide causes the fragmentation of mitochondria in SKOV-3 cells. **(a)** Representative confocal fluorescence images of mitochondria in SKOV-3 ovarian cancer cells treated with different concentrations of the 3D-NAF-1^44-67-6K^ peptide. **(b)** Quantitative analysis of mitochondrial length at different concentrations of the peptide. Data is shown as mean ± SD of 25 cells in 8 fields images calculated from 3 independent experiments. *\*\*\*P < 0.001*, *\*\*\*\*P < 0.0001*, by t-test.

**Fig. 4.**
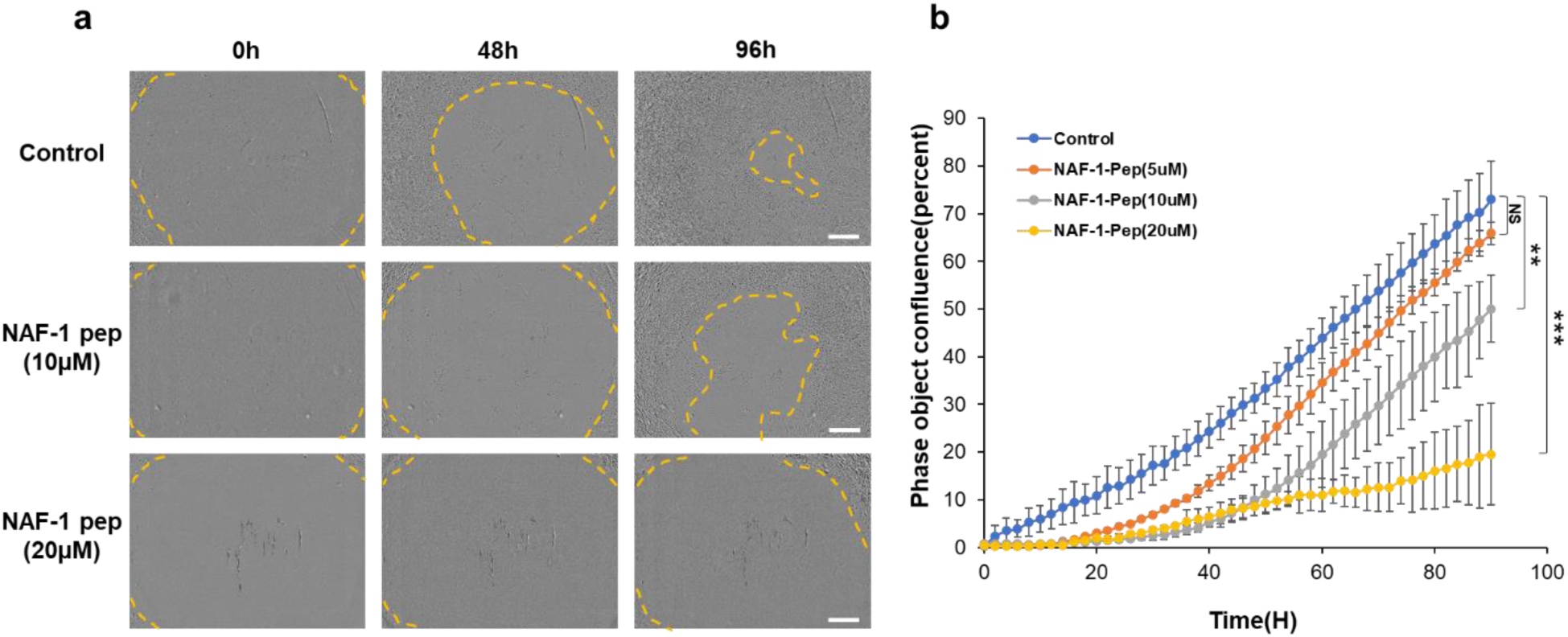
The effect of 3D-NAF-1^44-67-6K^ peptides on the migration of SKOV-3 cells. **(a)** Time-lapse high-definition phase contrast images of control untreated (upper panels) and 3D-NAF-1^44-67-6K^ treated (10μM, middle panels, 20μM, bottom panels) SKOV-3 cells at 0, 48, and 96 hours after culturing. The dotted lines define the area without cells. Scale bars, 200μm. **(b)** Quantification of cell migration during 96 hours of untreated (blue) and treated with 3D-NAF-1^44-67-6K^ peptides, 5μM (orange), 10μM (grey), 20μM (yellow). Results represent the mean of four measurements of cell migration, obtained in 3 independent experiments (n = 12). *\*\*P < 0.01, *** P < 0.001*. The corresponding p-values obtained by t-test.

### Treatment with the 3D-NAF-1^44-67-6K^ peptide suppresses the growth of ovarian cancer tumours in a human xenograft mice model

To study the effect of the peptide on ovarian cancer tumours, SKOV-3 cells were used to generate xenograft tumours in female mice. Control and xenograft mice carrying SKOV-3-derived tumours were then treated with the peptide (0.5 mg/kg) twice a week via either injections intra venously (IV) or subcutaneously (SC), while the control xenograft-carrying SKOV-3 mice were injected with saline. Following three weeks of treatment, the size of tumours in the 3D-NAF-1^44-67-6K^-treated mice significantly decreased compared to control (Fig. 5a). Based on mice weight, the treatment with the peptide appeared nontoxic, as it did not affect weight gain during the experiment (Fig. 5b).

**Fig. 5.**
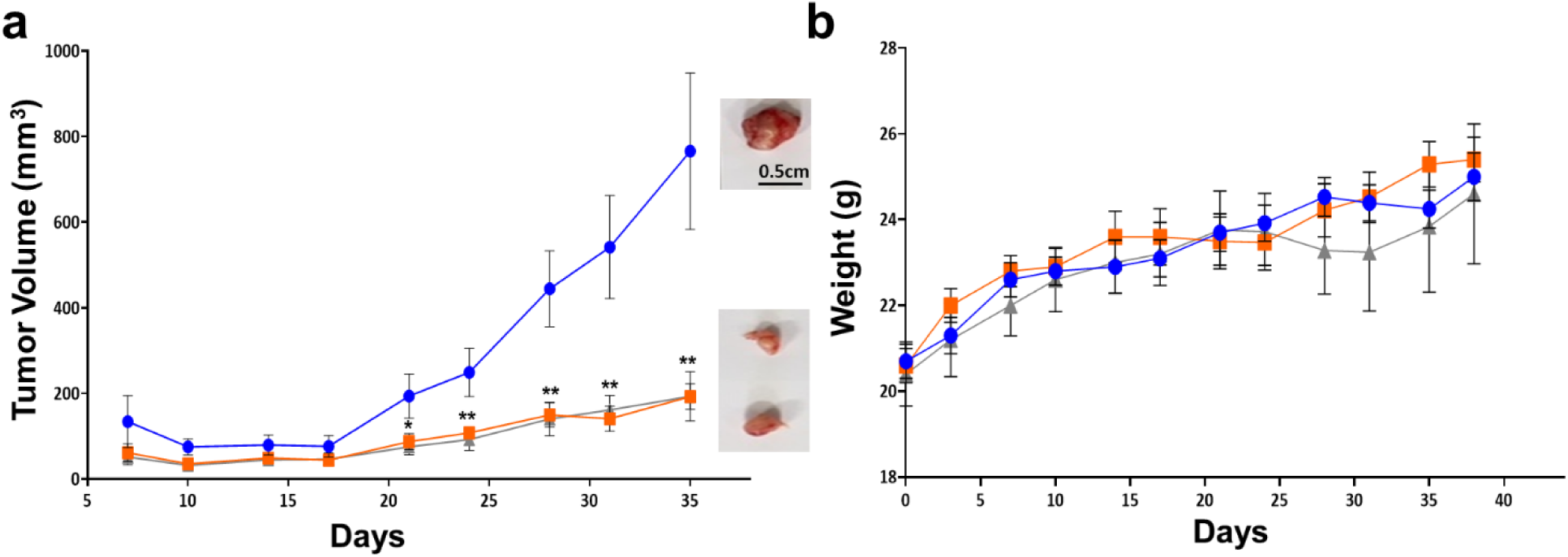
The impact of 3D-NAF-1^44-67-6K^ on primary xenograft SKOV-3 ovarian cancer tumours. **(a)** Tumour volume (mm^3^) of three mice groups: control (saline; blue), intravenously (IV; orange), and subcutaneously (SC; grey) injected groups. Representative tumours images are shown on right. **(b)** Changes in mice weight of the three groups over the treatment duration. 0.5 mg/Kg of 3D-NAF-1^44-67-6K^ peptide was used in each peptide treatment, injections were applied 6 times over the course of 3 weeks. All results are presented as mean ± SEM. Significance was determined using T-test; *\*P<0.1, **P<0.01*.

The toxicity of the 3D-NAF-1^44-67-6K^ peptide was further assessed by a chronic toxicity study, using male and female mice. Mice were injected with saline (control), or 3D-NAF-1^44-67-6K^, using two concentrations: *i*) the concentration used for the treatment described in Fig. 5 (0.5 mg/kg) and *ii*) a 10-fold higher concentration of (5 mg/kg). At the end of the study, complete blood count (CBC) changes were determined. These showed no difference between treated and un-treated mice groups (Supplementary table 2). In addition, organs characteristics of mice were evaluated for their weight, as well as histopathological changes following sectioning and H&E staining. The results of both treated groups were identical to that of control mice, further demonstrating that the peptide was non-toxic to healthy cells and organs (Supplementary Fig. 8). These findings demonstrate the low and potentially non-toxicity of the peptide toward healthy mice cells *in-vivo* (Supplementary Table 2 and Supplementary Fig. 8).

### Treatment with the 3D-NAF-1^44-67-6K^ peptide decreases the metastatic activity of SKOV-3 ovarian cancer cells *in vivo*

Ovarian cancer cells are known to have high mobility and metastatic activity^45^. The findings that the 3D-NAF-1^44-67-6K^ peptide significantly decreases the migration of SKOV-3 cells *in vitro* (Fig. 4) encouraged us to study the effects of the 3D-NAF-1^44-67-6K^ peptide on the metastatic activity of SKOV-3 cancer cells in mice. For this purpose, we allowed SKOV-3 cells to metastasize following their initial IV injection resulting in lung metastasis/nodules. We then compared the effects of treatment with the 3D-NAF-1^44-67-6K^ peptide to that of Taxol^46, 47, 48^, one of the primary chemotherapy drugs used to treat OC. The peptide (0.5 mg/Kg) was found to have a similar effect to that of Taxol at (20 mg/Kg), suppressing ovarian cancer metastasis in lungs (Fig. 6a). In addition, the peptide was able to suppress metastasis into the colon of mice (Fig. 6b).

**Fig. 6.**
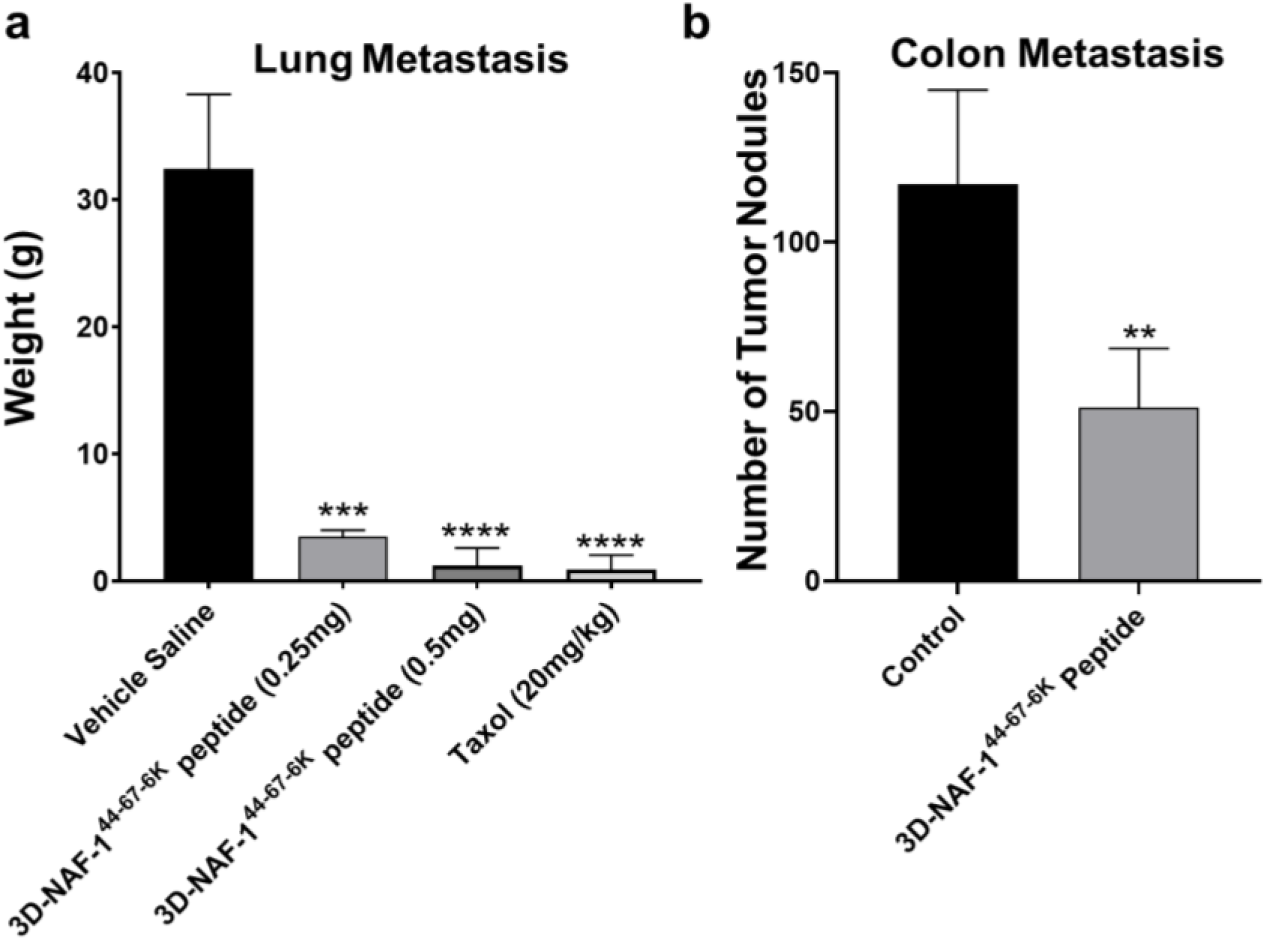
3D-NAF-1^44-67-6K^ suppresses metastatic growth of xenograft SKOV-3 ovarian cancer. **(a)** Decrease in tumour nodule weight (gr) in lungs of mice injected IV with SKOV-3 cells and treated with 3D-NAF-1^44-67-6K^ peptide at either 0.25 or 0.5 mg/Kg per mouse. Injections were performed twice a week intravenously (IV). Taxol 20 mg/Kg (once a week) IV was used as positive control. **(b)** Decrease in colon nodule metastasis (Nodules number) following treatment with Saline (control) or 0.5mg/Kg 3D-NAF-1^44-67-6K^ peptide. All results were presented as mean ± SEM. Significance was determined using T-test; *\*\*P<0.01, ***P<0.001, ****P<0.0001*.

### Broad-spectrum anti-cancer activity of the 3D-NAF-1^44-67-6K^ peptide

As 3D-NAF-1^44-67-6K^ demonstrated anti-cancer activity towards breast and ovarian cancer cells (Figs. 1-6; Supplementary Fig. 1), we studied its killing effect on several other cancer cells including acute myeloid leukaemia (MOLM-14), chronic myelogenous leukaemia (K562), pancreas cancer (PANC-1), brain glioblastoma cancer cells (U87MG), and human malignant melanoma (A375). As shown in Fig. 7, at a low concentration of 5μM, the peptide decreased the viability of all cancer cells, with the most toxic effects towards the leukaemia cancer cells. At the higher concentrations of 25 μM, the peptide dramatically decreased the viability of all tested cancer cells lines. Taken together with our findings with breast and ovarian cancers, these findings suggest that the 3D-NAF-1^44-67-6K^ peptide could be used against a wide spectrum of cancers.

**Fig. 7.**
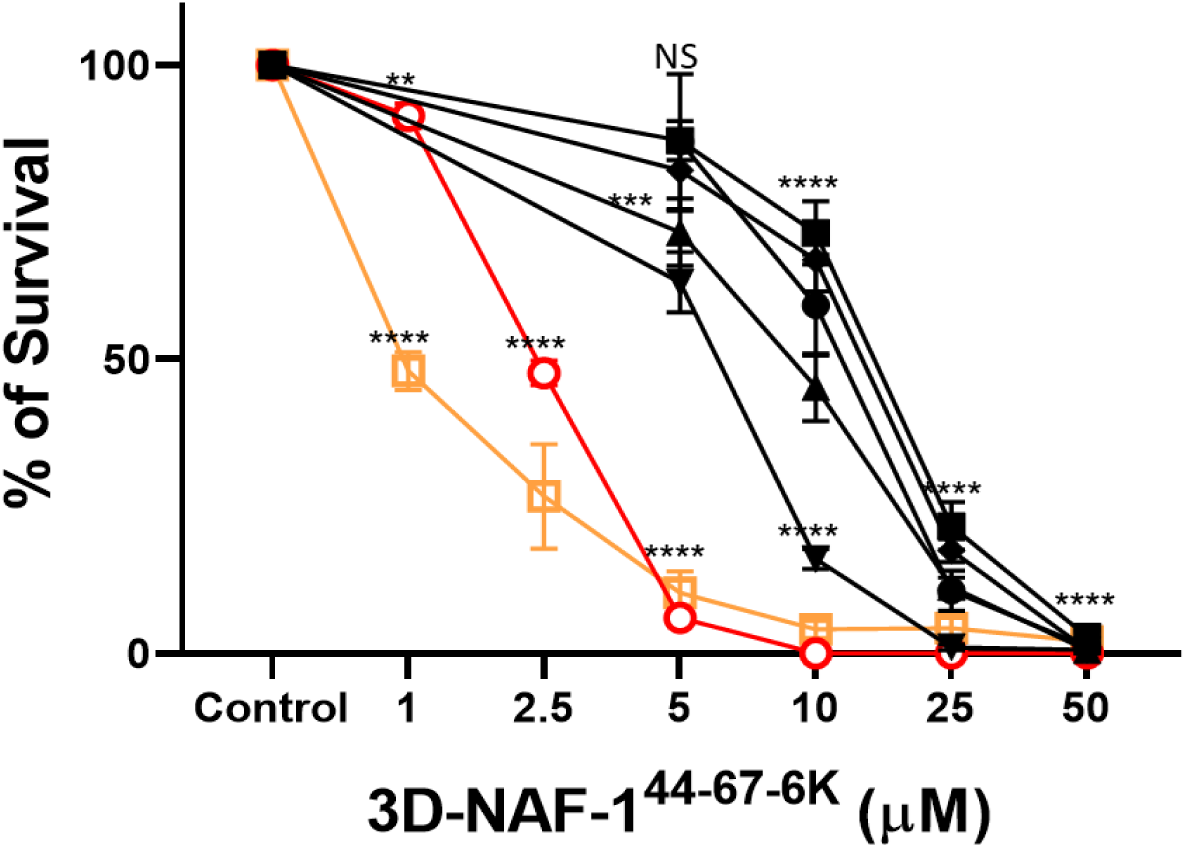
Cytotoxicity of 3D-NAF-1^44-67-6K^ towards different cancer cell lines. Cell survival was measured using the presto-blue assay following 3 hours incubation with the peptide at the indicated concentrations.SKOV-3, ovarian cancer (black circle), U87MG, glioblastoma (black rectangle), A375, human malignant melanoma (black up-point triangle), MDA-MB-231, breast cancer (black down-point triangle), PANC-1, pancreas cancer (black diamond), MOLM-14, acute myeloid leukaemia (red circle), and K562, chronic myelogenous leukaemia (yellow rectangle) were treated or untreated with the peptide and assayed for viability and survival. Data are shown as mean ± SD from three independent experiments. *\*\* P<0.01, *** P<0.001, **** P<0.0001* by t-test.

## Discussion

We previously reported on a peptide derived from the CISD2/NAF-1 protein, 3D-NAF-1^44-67^, which selectively permeates breast cancer cells leading to their death, without affecting control non-cancerous breast cancer cells^31^. Here, we introduce an improved variant of this peptide which has been engineered to contain a longer C-terminus harbouring six positively charged lysine residues (3D-NAF-1^44-67-6K^). Our findings indicate that the positively charged 6K tail, added to the peptide, was only partially responsible for its increased toxicity, as adding six positively charged arginine residues (3D-NAF-1^44-67-6R^) resulted in significantly lower toxicity than that of 3D-NAF-1^44-67-6K^ (Supplementary Fig. 1&2). Our simulation results suggest that, compared to the original 3D-NAF-1^44-67^ peptide, the improved toxicity or the 3D-NAF-1^44-67-6K^ is due to the fact that the 6 Lysine tail pulls the centre-of-mass of the peptide closer to the cancer cell PM (Supplementary Fig. 3), and that this pull results in an improved permeation of the 3D-NAF-1^44-67-6K^.

The improved anticancer activity of 3D-NAF-1^44-67-6K^ prompted us to test its cytotoxic activity against other cancers including OC. Our study reveals that 3D-NAF-1^44-67-6K^ is toxic toward both OVCAR-3 and SKOV-3 OC cell lines, with a slightly higher toxicity towards OVCAR-3 (Fig. 1). 3D-NAF-1^44-67-6K^ was further found to suppress primary tumour growth of SKOV-3 xenograft tumours (Fig. 5), and more importantly, to significantly decrease metastatic growth of SKOV-3 cells in mice, in both lungs and colon (Fig. 6). The treatment of mice with the 3D-NAF-1^44-67-6K^ peptide did not cause weight loss, indicating that the peptide is not toxic (Fig 5 and supplementary Fig. 8). This finding was further supported by a chronic toxicity study that examined blood and organs from control mice mice injected with 0.5mg/Kg that showed no apparent cytotoxic effects with even 10-fold higher concentrations (5mg/Kg) of the peptide in mice (Fig. 5 and Supplementary Fig. 8). Further studies are however required to test for toxic and/or other side effects of the peptide in animals and patients.

3D-NAF-1^44-67-6K^ was shown to permeate the PM of OC cells and target their mitochondria (Fig. 2). The peptide induced mitochondrial fission that caused a significant decrease in mitochondrial length and membrane potential (Fig. 3). This was accompanied by a dramatic decrease in SKOV-3 cell’s migration activity and viability (Fig. 1 and Fig. 4 and Supplementary Fig. 4). The mitochondrial targeting properties of 3D-NAF-1^44-67-6K^ could be a reflection of its biological origin (*i.e.,* the subcellular localization of the protein its derived from CISD2/NAF-1). CISD2/NAF-1 is a transmembrane protein, localized to the mitochondrial outer membrane and the endoplasmic reticulum (ER), and their connecting membranes (MAM), and the 3D-NAF-1^44-67-6K^ peptide is derived from part of the transmembrane and flexible/unstructured domains of CISD2/NAF-1 ^31^.

Although other anticancer peptides were shown to suppress metastatic cancer growth^26, 28, 29, 30^, only few showed such a broad-spectrum activity towards many different cancers (Figure 7). The polyarginine-p53 cell penetrating peptide (CPP; R11-p53) was for example shown to selectively penetrate bladder cancer cells and was able to decrease both tumour growth and metastatic growth in four different bladder cancer cell lines^32, 49^. Both our peptide and R11-p53C showed therefore selectivity in permeating cancer cells, however, while R11-p53C specifically targeted blabber cells, the NAF-1 derived peptide was able to target many different types of cancer cells (Fig. 7). The selective mode of function of the 3D-NAF-1^44-67-6K^ peptide towards cancer cells of many types of solid tumours (breast, ovarian, melanoma and glioblastoma) and against different blood malignancies (Fig. 7), suggests that this peptide might act as a more universal cancer-specific drug, exploiting the common features of cancer cell PMs^31, 50,51, 52^ (Supplementary table 1 and Supplementary Fig. 3).

Taken together, our findings indicate that the 3D-NAF-1^44-67-6K^ peptide selectively permeates OC cells and causes significant mitochondrial fission damage, leading to OC cell death (Fig. 1-3). As indicated by weight gain and a chronic toxicity study in mice, the 3D-NAF-1^44-67-6K^ peptide appears to be non-toxic to mice. In addition, we show that the 3D-NAF-1^44-67-6K^ peptide can target both primary and metastatic OC tumours (Fig. 5-6), as well as multiple other cancers (Fig. 7). These finding suggest that the 3D-NAF-1^44-67-6K^ peptide could be an ideal candidate for clinical trials in ovarian cancer patients, and that it could also be used to target other cancer types. The findings that the 3D-NAF-1^44-67-6K^ peptide has no apparent toxicity toward non-cancerous cells further suggest that it could be used in combination therapy treatments, together with other anticancer drugs. We hope that our study will facilitate the development of much needed highly efficient OC treatments.

## Methods

### Xenograft model

Female Nude CD1 mice were used for studying OC xenograft tumour cell progression of SKOV-3 cells, injected subcutaneously. The experiment was approved by the Authority for Biological and Biomedical Models at the Hebrew University, ethical number is NS-13-13911-4. SKOV-3 of 5X10^6^ cells/mouse were injected subcutaneously to the flank of each mouse. Each mice group contained 5-10 mice. Mice were injected (2 times/week) for a total of 6 injections, either subcutaneously (SC) or intravenously (IV), with 0.5g/Kg (∼50 µM/mouse) of the 3D-NAF-1^44-67-6K^ peptide, or injected with Saline for the control group. Tumour volume (mm^3^) was measured every 3 days before and following injections as described in^31^. Toxicity of the treatment was evaluated by measuring mice weight change (g) that was measured twice a week^33^. All results were presented as mean ± SEM.

### Chronic Toxicity Investigation

Control male and female NOD/SCID mice groups (4-5 mice per group), were injected intraperitonially (IP) with 3D-NAF-1^44-67-6K^ peptide using a concentration of 0.5mg/Kg or 5mg/Kg, while a control group was injected with Saline; total volume of each injection was 100µl. Mice were injected 3 times a week for 4 weeks, the weight of the mice was measured and showed no difference between the groups during the whole experiment (Supplementary Fig. 8.a). At the end of the experiment, mice were terminated and blood samples were collected from all groups, using EDTA-CBC tubes, and analysed using the ADVIA® 2120i System (Siemens, Germany), at the laboratory of Haematology at the Hadassah medical center ^53^. In addition, organs (heart, kidneys, liver, lungs and spleen) were collected, weighed and subjected to histopathological investigation by staining with hematoxylin and eosin (H&E), at the histopathological laboratory of the Authority for biological and biomedical models, at the Hebrew university of Jerusalem, as described previously^53^.

### Metastasis model

Female mice of Nude CD1 were injected (intravenously) into the tail vein with 1x10^6^ cells/mouse of SKOV-3 cells prepared in PBS (200μl/mouse). Ethical approval number NS-19-15989-5. Treatment of animals started four weeks after cancer injection, with 0.25 or 0.5 mg/Kg of 3D-NAF-1^44-67-6K^ peptide for each mouse (as described above); as a control either vehicle saline was injected, or the well-known OC chemotherapy treatment Taxol (20 mg/Kg) was used as positive, each group contained 10 mice. Lung and colon metastasis were observed in the Xenograft nude mice as described above.

### Cell cultures

Ovarian cancer cells (SKOV-3; ATCC, HTB-77^TM^)^54^, human malignant melanoma (A375), were grown in 5% CO_2_ DMEM – high glucose (SIGMA, D5796) medium supplemented with 10% FCS, L-glutamine, and antibiotics (Biological Industries). Malignant epithelial breast cells (MDA-MB-231) were grown in 5% CO_2_ RPMI medium 1640 supplemented with 10% FCS, L-glutamine, and antibiotics (Biological Industries). Acute myeloid leukaemia (MOLM-14), and chronic myelogenous leukaemia (K562) were grown in 5% CO_2_ DMEM – high glucose (SIGMA, D5796) medium supplemented with 10% FCS, L-glutamine, and penicillin-streptomycin antibiotics (Biological Industries). Epithelial normal breast cells (MCF-10A) were grown in complete growth medium consisting of 1: 1 mixture of Dulbecco’s modified Eagle’s medium and Ham’s F12 medium supplemented with horse serum (5%), epidermal growth factor (20 ng ml_1), cholera toxin (CT, 0.1 mg mg^-1^), insulin (10 mg ml^-1^), hydrocortisone (500 ng ml^-1^), and penicillin/streptomycin (1 unit per ml). Pancreas cancer (PANC-1), glioblastoma (U87MG) were grown in 5% CO_2_ DMEM – high glucose (SIGMA, D5796) medium supplemented with 10% FCS, L-glutamine, and penicillin-streptomycin antibiotics (Biological Industries). Cells were plated one day prior to the experiments on 96-well plates or 4-Chamber Glass Bottom Dish for cell viability or microscopic measurements.

### Confocal microscopy measurements

Cells 2 × 10^5^, were planted in μ-slide 4-well with glass bottom one day prior to the experiment. For cell permeation experiments, cells were incubated with Fl-3D-NAF-1^44-67-6K^ (10μM), Fl (10 μM) alone, Cy-3D-NAF-1^44-67-6K^ (10μM) or Cy5 alone (10μM), for 6h. Nuclei were stained with Hoechst 33342 and mitochondria with Rhodamine 800 or mito-green. Fluorescence was followed with confocal microscopy (Olympus FV3000 confocal laser-scanning microscope). For mitochondrial morphology experiment, cells were incubated at different concentrations of 3D-NAF-1^44-67-6K^. mitochondria were stained with RPA, red fluorescence was captured with the confocal microscopy, and all images were analysed with image J^55^.

### Mitochondrial length measurement

To measure mitochondrial length and number, binary images were obtained through the thresholding function of image J to select the objects to be skeletonized by MiNA (Mitochondrial Network Analysis) program which yields mitochondrial network parameters such as the mean branch length, number of individuals and networks, and mitochondrial footprint. The skeleton length and number formed with MiNA are measured for each frame of selected ROI. Quantification of mitochondrial length and number was performed on the samples as describe in details (Supplementary Fig. 7). The detailed steps of MiNA are described in ref.^56^.

### SKOV-3 cells viability assay

SKOV-3 cells were plated in a 96-well plate at a density of 12 x 10^3^ and were allowed to culture for one day. 3D-NAF-1^44-67-6K^ was added to the cells at concentrations of 0, 5, 10, 15, 20, and 25μM and cells were incubated with the peptide for up to 48 hours. Cell viability was measured at three time points: 6, 24, and 48 hours. Cell viability was determined by the fluorescent redox probe, Presto-blue^TM57^ (Invitrogen™, A13261). Cells were incubated with Presto-blue for 1 hour and Presto-blue fluorescence was measured at 37°C (λ_ex_=560 nm, λ_em_=590nm) by a plate reader (Tecan Safire), every group contains four repeats, results represented as mean ± SD.

### OVCAR-3 cells viability assay

OVCAR-3 cells were plated in a 96-well plate at a density of 15 x 10^5^ and were allowed to culture for one day. 3D-NAF-1^44-67-6K^ was added to the cells at concentrations of 0, 2.5, 5, 10, and 20μM and cells were incubated with the peptide for up to 24 hours. Cell viability was measured at two time points: 3 and 24 hours. Cell viability was determined by the fluorescent probe Propidium iodide (Sigma™, 25535-16-4). Cells were incubated with Propidium iodide for 15 minutes and Propidium iodide fluorescence in cells was observed at 37°C (λ_ex_=525 nm, λ_em_=615nm) by Incucyte®. Every group contains four repeats, results represented as mean ± SD.

### Cytotoxicity of ovarian cancer cells 3D-spheroids

SKOV-3 cells were plated in a 96-well plate at a density of 2000 cells/well and were allowed to culture for 3 days to form 3D-spheroids. 3D-NAF-1^44-67-6K^ was added to the spheroids at concentrations of 0, 15, 25, and 50μM and IncuCyte® Red cytotoxicity reagent (Essen Bioscience Cat #4632) was used to probe for cell death. Spheroid images were recorded every hour for 15 hours. The images and the resulted red colour, which indicates cell death, were analysed with the IncuCyte® Zoom system (Essen Bioscience). Every group contains three repeats, results represented as mean ± SD.

### Determination of IC50

SKOV-3 cells or OVCAR-3 cells were planted at a density of 12 × 10^3^ cells/well and 15 × 10^3^ in a 96 well plate, respectively, and treated with the 3D-NAF-1^44-67-6K^ peptide at the following series of concentrations (0.65, 1.25, 2.5, 5, 10, 20, 40, 80, 120, and 140 µM) for 3 hours. % of cell death was calculated in reference to cells without the peptide. The IC50 values were calculated by fitting the plot to the Hill equation: Y = B + ((T - B)/(1 + 10(log^(EC50)-X)^ ^×^ ^Hill^ ^slope^)) T and B: top and bottom plateaus in the Y axis; IC50: the X value that gives the half Y value between top and bottom Y plateaus. Hill slop is the curve slope. The experiment contains four repeats, results represented as mean ± SD.

### Cell migration and invasion assay

SKOV-3 cells were plated in a 96-well plate with the round glass cover slip in the center of the well at a density of 2 x 10^4^ cells/well and were allowed to culture for 1 days. Following day, cells were treated with 3D-NAF-1^44-67-6K^ peptides (0μM, 10μM, 20μM) after the removal of the round glass cover slide. Cell phase contrast images were recorded every three hours for 96 hours. The phase contrast images were analysed with the IncuCyte® Zoom system (Essen Bioscience).

## Acknowledgements (optional)

This work was supported by the BSF grant number 2020094 (to R.N. and A.F. and R.E.), National Institute of Health grant GM111364 (to R.E. and R.M.). We thank The Charles E. Smith Family and Prof. Joel Elkes Laboratory for Collaborative Research.

## Author contributions

Conceptualization: RN, AF, RM, RE, EP. Experimental design: RN, AF, RM. Experimental data collection and analysis: EN, Y-SS, OK, M.DY. All authors wrote the manuscripts.

## Supplemental Material

### 1. Improved cytotoxicity of 3D-NAF-1^44-67-6K^

3D-NAF-1^44-67-6K^ has higher cytotoxicity towards breast cancer cells than the 3D-NAF-1^44-67^ peptide, while remaining non-toxic towards healthy cells.

**Supplementary Figure 1.**
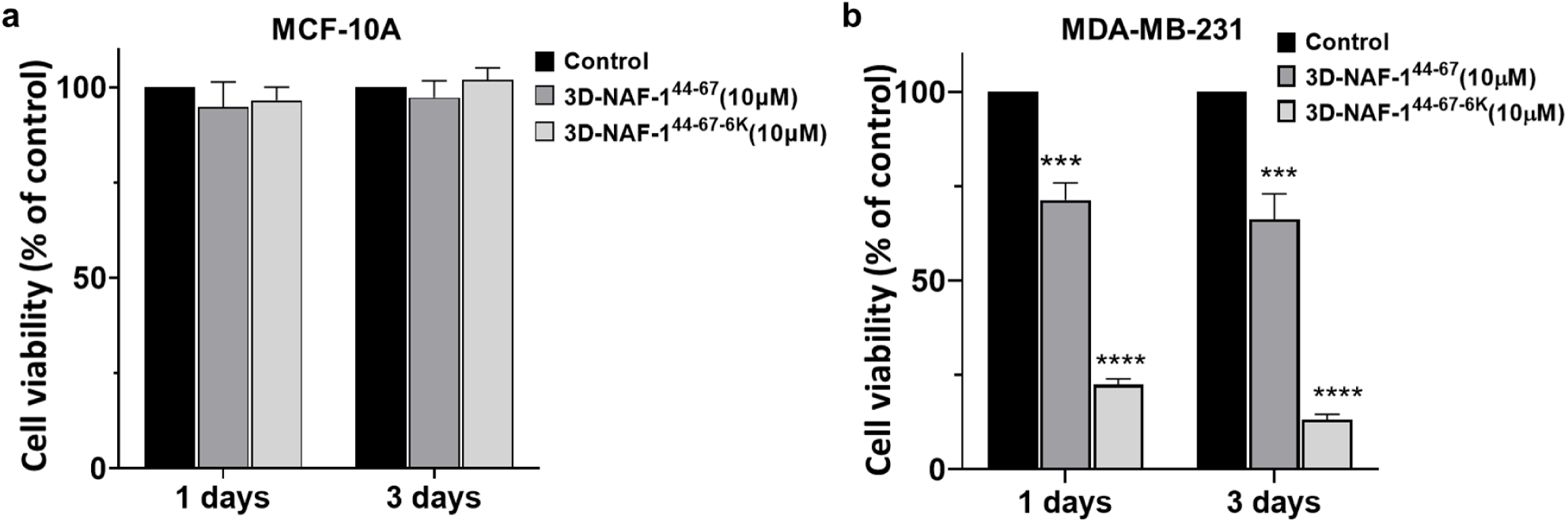
Cytotoxicity of the 3D-NAF-1^44-67^ and 3D-NAF-1^44-67-6K^ peptides towards control MCF-10A cells (a) and malignant MDA-MB-231 cells. **(b)**. Cells were plated in 96 well plates at the density of 15,000 per well before the experiment. 10µM of the 3D-NAF-1^44-67^ and 3D-NAF-1^44-67-6K^ peptides was added to cells and incubated for 1 or 3 days. Cell viability was assessed using Presto-Blue after 1- and 3-days incubation. Statistic data were obtained against the control (without peptides) *\*\*\*P < 0.001*, *\*\*\*\*P < 0.0001* by t-test.

### 2. 3D-NAF-1-^44-67-6K^ *vs.* 3D-NAF-1-^44-67-6R^

The higher cytotoxic activity of the 3D-NAF-1-^44-67-6K^ peptide is not a result from just the addition of the six positively charged residues. As Supplementary Figure 2 shows addition of 6 positively charged Arginines did not have the same cytotoxic effect.

**Supplementary Figure 2.**
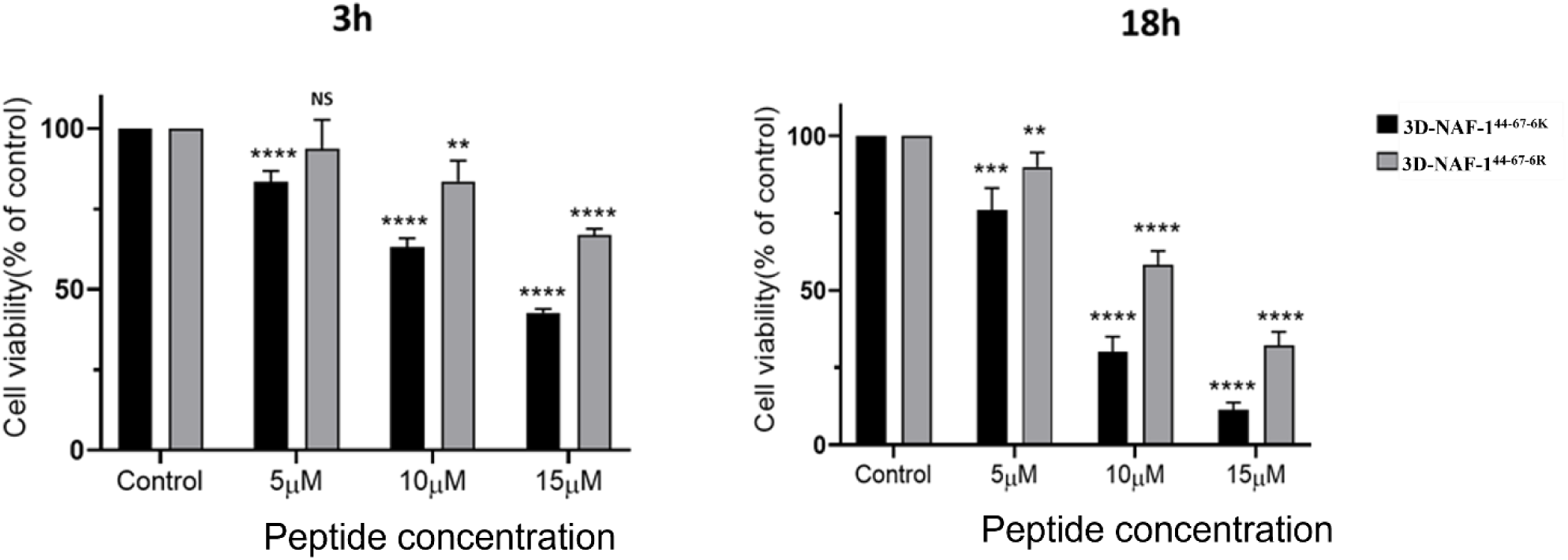
The cytotoxic effect of 3D-NAF-1^44-67-6K^ on MDA-MB-231 cells is higher than that of 3D-NAF-1^44-67-6R^. Cells were seeded in a 96-well plate at a density of 1.2 x 10^4^ and allowed to culture for overnight 3D-NAF-1^44-67-6K^ or 3D-NAF-1^44-67-6R^ peptides were incubated with cells for 3 and 18 hours at different concentrations. The black bars indicate the cytotoxic effect of 3D-NAF-1^44-67-6K^, while the grey bars show the cytotoxicity of 3D-NAF-1^44-67-6R^. Cell viability was determined using the fluorescent redox probe, Presto-blue^TM^. Cells are incubated with Presto-blue for 1 hour and Presto-blue fluorescence was measured at 37°C (λ_ex_=560 nm, λ_em_=590nm) by a plate reader (Tecan Safire). The results indicate that the 3D-NAF-1-6K is more toxic to malignant cells than the 3D-NAF-1-6R SO the addition of 6K is more than just 6 positive charges. NS = none significant. Statistic data were obtained against the control (without peptides) *\*\*P < 0.01***P < 0.001*, *\*\*\*\*P < 0.0001* by t-test.

As stated in paper, in order to better understand the higher cytotoxicity of the 3D-NAF-1^44-67-6K^ we used computational simulation to analyze what is the effect of the 6-Lysine addition on the permeation to malignant vs. normal membranes.

### 3. Permeation simulation

#### Early permeation simulation of 3D-NAF-1^44-67^ and 3D-NAF-1^44-67-6K^ into healthy/malignant cells

The permeation of both 3D-NAF-1^44-67^ and 3D-NAF-1^44-67-6K^ into healthy and epithelial breast cancer cells membrane were simulated computationally (Supplementary Fig. 3).

During the 1.5 μs trajectories, NAF-1^44-67^ moves closer to the cancer membrane surface but remained farther away from the normal membrane. The top panels of Supplementary Fig. 3.a show representative molecular views of those general tendencies. Similar views for NAF-1^44-67-6K^ show a larger number of surface membrane interactions for this peptide on both membranes (bottom panels of a). Supplementary Fig. 3.b quantifies the different degrees of approach and insertion of the center of mass of the peptides in the two membrane models. For the normal membrane (red lines), the center of mass of NAF-1^44-67^ is far from the membrane surface (in the membrane the phosphates of the phospholipids molecules are located at ∼1.8 nm from the membrane center), while the center of mass of NAF-1^44-67-6K^ is closer to the membrane/water interface. For the cancer membrane (black lines), the center of mass of NAF-1^44-67^ is closer to the surface so some parts of the peptide interact with the lipids. The closest approach of the center of mass of the peptide to the membrane occurs when NAF-1^44-67-6K^ interacts with the cancer membrane. The association and insertion of both peptides to the cancer membrane points out to the influence of attractive electrostatics interactions during the approach of the peptide to the more negatively charged membrane.

Which part of the peptide interacts more with the phospholipids during these early insertion events? Supplementary Fig. 3.c shows the accumulated number of interactions of the N-terminal part of the peptide (that contains mostly hydrophobic residues) and the C-terminal part (that contains all the positively charged residues) with the phospholipid heads. For NAF-1^44-67-6K^ the hydrophobic part of the peptide has a larger number of interactions with the phospholipid heads than NAF-1^44-67^ reflecting a deeper insertion of the N-terminal part of NAF-1^44-67-6K^ (see also the panels in A). The charged part of both peptides interacts similarly with the cancer membrane model (the right-side panels of Supplementary Fig. 2.a show that the C-terminal for both peptides “lie down” on the surface of the cancer membrane models). For the normal membrane, the number of interactions with the C-terminal of the peptides is smaller (see also left-side panels of Supplementary Fig. 2.a). When comparing the C-terminal interactions of NAF-1^44-67-6K^ with both model membranes headgroups we observed almost twice the number of interactions with the cancer membrane than the normal membrane. That expanding number of interactions with the charge part of the peptide could facilitate the full permeation event in the cancer membrane.

**Supplementary Table 1:**
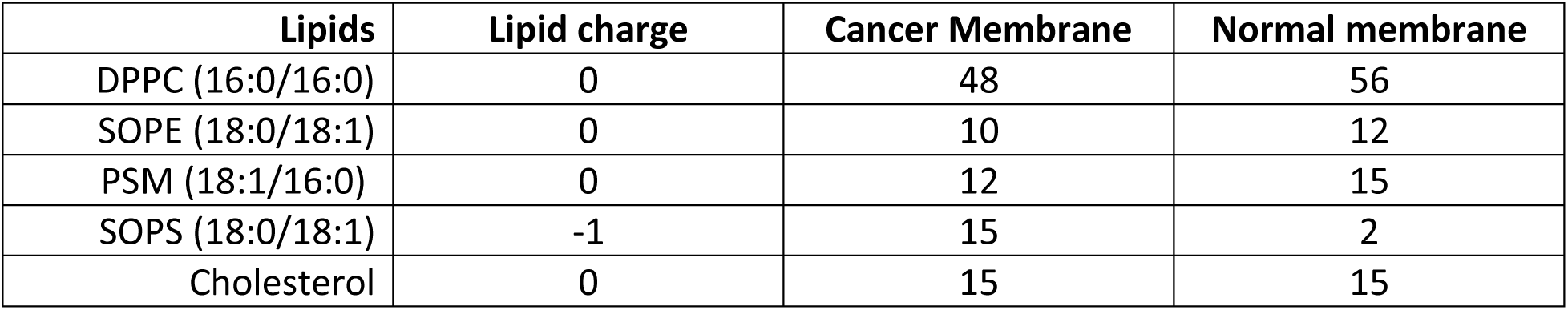
Lipid compositions for the cancer and normal membrane models used in the simulations. The Lipids column provides the common abbreviations for the phospholipids and the size and amount of unsaturation of their two tails. We built symmetric membrane models (both layers of the membranes have the same composition). Experimentally, it has been found that in most cancers the plasma membranes have a larger number of negatively charged PS on its external layer^1, 2^ than in normal membrane. We modeled that variation in PS numbers in our membranes.

**Supplementary Figure 3:**
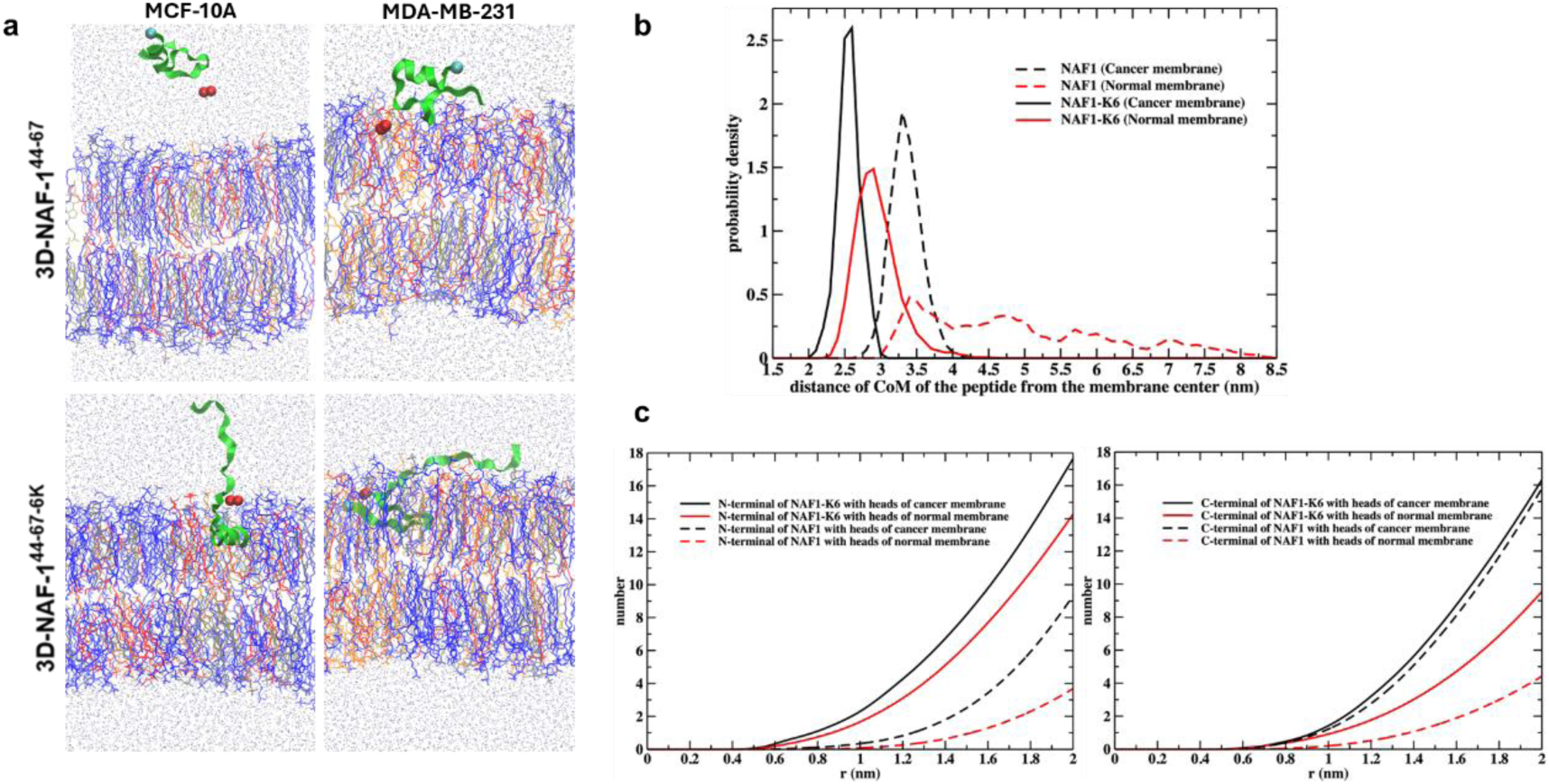
Early permeation simulation of 3D-NAF-1^44-67^ and 3D-NAF-1^44-67-6K^ into healthy/malignant cells. **(a)** Representative views of the peptide interacting with the model membranes. The top two panels are for NAF-1^44-67^ and the bottom panels for NAF-1^44-67-6K^, the left side shows the normal membrane (modeling the MCF-10 cell line) and the right side shows the cancer membrane (MDA-MB-23). The lipids are shown with lines of different colors: DPPC (blue), SOPE (red), SOPS (orange), PSM (gray), and cholesterol (tan). The backbone of the peptide is shown with green ribbons. The locations of some atoms of the peptide are displayed as spheres: the Cα of F1 (cyan) and the terminal nitrogen atoms containing the positive charge of R14 (red). These molecular representations were generated using VMD^3^. **(b)** Probability density to find the center of mass of the peptide (NAF-1^44-67^ or NAF-1^44-67-6K^) at different separations from the membrane center.**(c)** Accumulated number of interactions between different phospholipid heads and the N-terminal half (left panel) or C-terminal half of the peptides up to a separation of 2 nm between the center of mass of the respective terminal and the heads of the phospholipids. For NAF-1^44-67^ and NAF-1^44-67-6K^ the N-terminal half consists of residues 1-13, and the C-terminal is comprised for the rest of the residues in the peptide sequences.

### 4. IC50 measurements

#### The 3D-NAF-1^44-67-6K^ is toxic towards SKOV-3 and OVCAR ovarian cancer cell lines

**Supplementary Figure 4.**
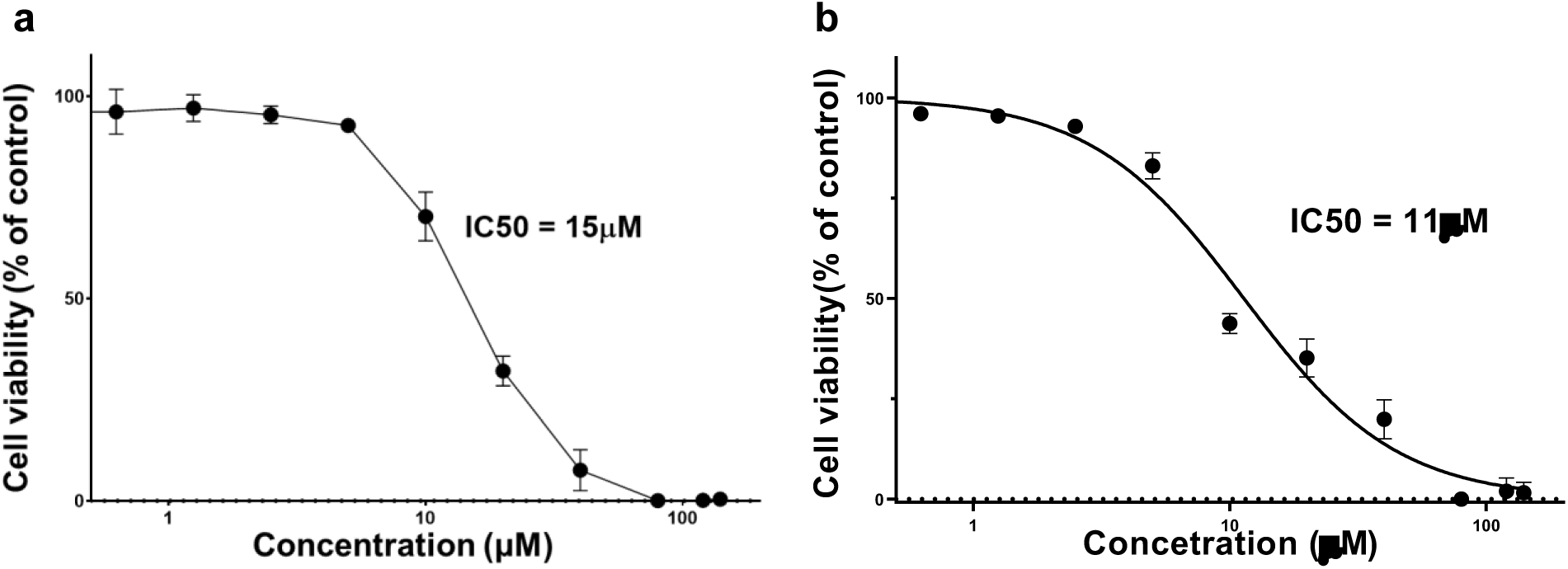
IC50 of 3D-NAF-1^44-67-6K^ towards (a) SKOV-3 cells and OVCAR-3 **(b).** IC50 value for inducing SKOV-3 or OVCAR-3 cell death by 3D-NAF-1^44-67-6K^ peptide at 3 hours of treatment. The IC50 value was found to be 15μM and 11μM for SKOV-3 and OVCAR-3, respectively. The details of IC50 determination are described in the method.

### 5. Permeation of fluorescent tagged 3D-NAF-1^44-67-6K^

#### 3D-NAF-1^44-67-6K^ permeate selectively into ovarian cancer cells

**Supplementary Figure 5.**
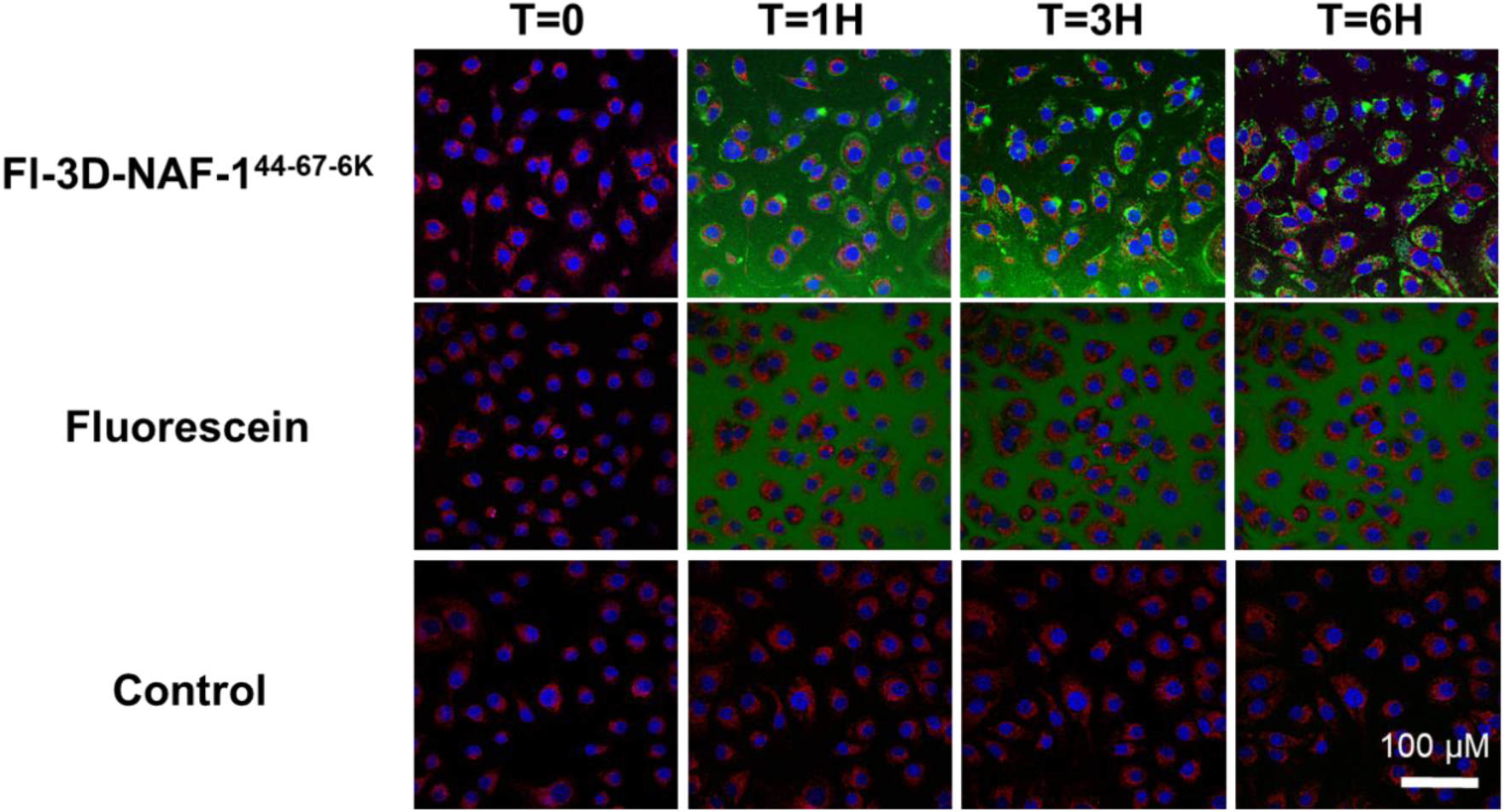
Permeation of Fl-3D-NAF-1^44-67-6K^ into ovarian cancer cells and targeting of mitochondria. Representative confocal fluorescence images of SKOV-3 ovarian cancer cells at different time points following treatment with and without Fl-3D-NAF-1^44-67-6K^ peptide and fluorescein alone. The quantitative graph for the permeation of Fl-3D-NAF-1^44-67-6K^ peptide into ovarian cancer cells is depicted in figure 2 of the main manuscript. T=0 measured before the addition of the Fluorescein or Fl-3D-NAF-1^44-67-6K^ peptide.

**Supplementary Figure 6.**
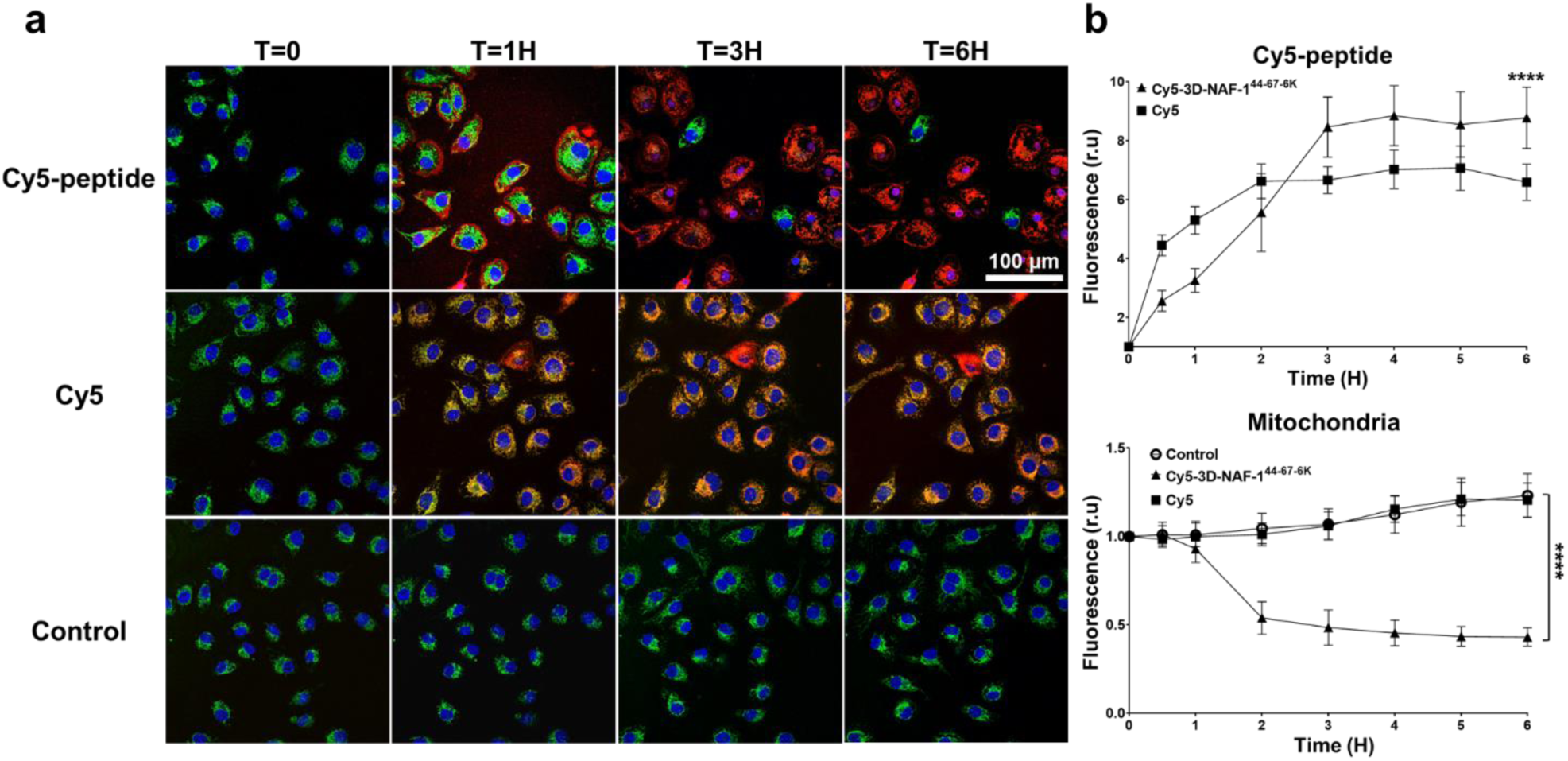
Permeation of Cy5-3D-NAF-1^44-67-6K^ into ovarian cancer cells and targeting of mitochondria. **(a)** Representative confocal fluorescence images of SKOV-3 ovarian cancer cells at different time points following treatment with and without Cy5-3D-NAF-1^44-67-6K^ peptide and Cy5 alone. T=0 measured before the addition of the Cy5 or Cy5-3D-NAF-1^44-67-6K^ peptide. **(b)** Quantitative analysis of Cy5-3D-NAF-1^44-67-6K^ permeation into the SKOV-3 cells (upper) and mitochondrial damage (bottom) as accessed by the decrease in the green fluorescence of mito-tracker due to the damage in the mitochondria. Data is shown as mean ± SD of 15 cells per field (10 fields in total) calculated for each time-point obtained from 3 independent experiments. *\*\*\*\* P < 0.0001* by t-test.

### 6. Mitochondrial fragmentation

#### 3D-NAF-1^44-67-6K^ causes mitochondrial fragmentation in ovarian cancer cells

**Supplementary Figure 7.**
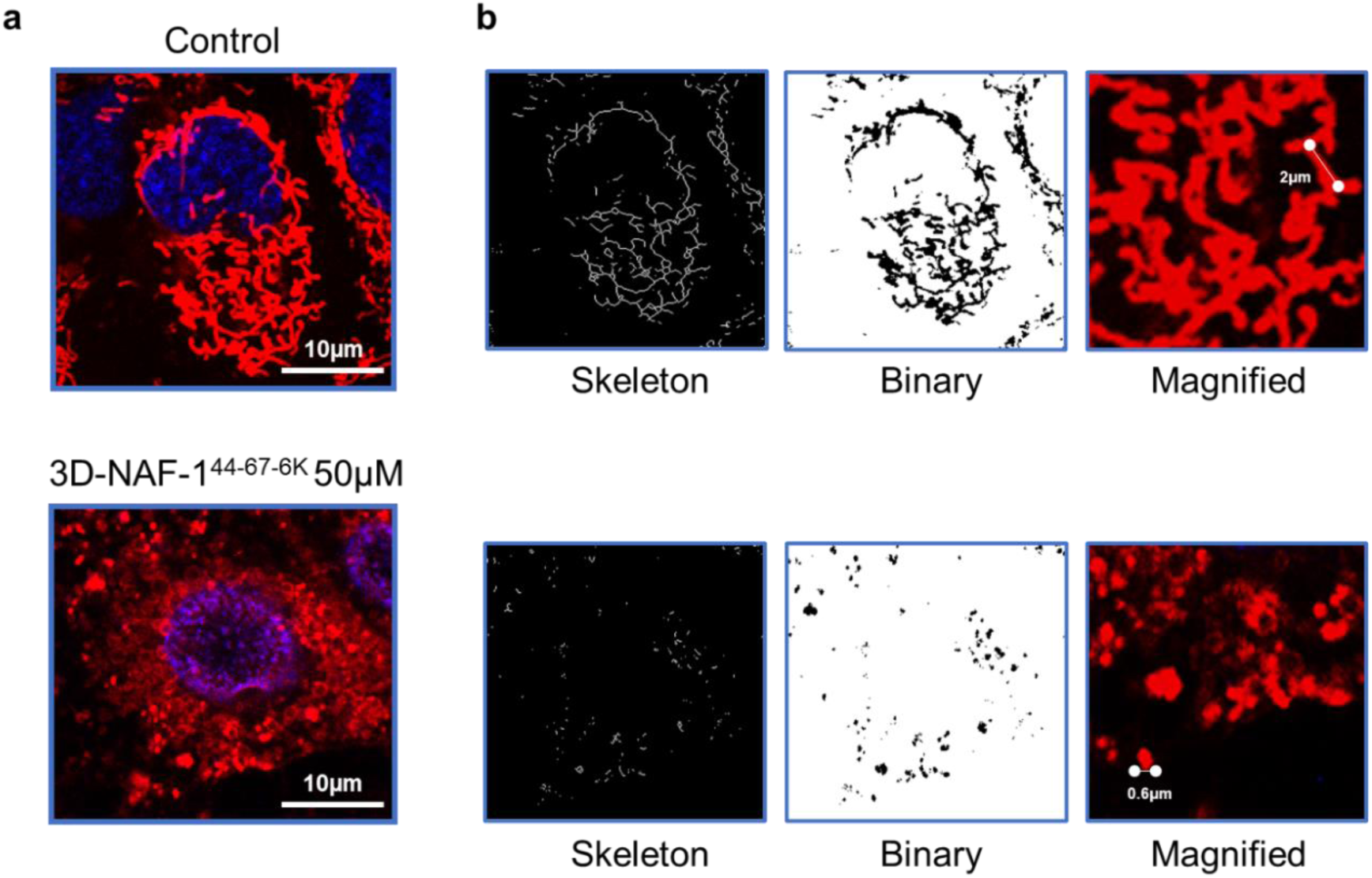
Measurement of mitochondrial length in SKOV-3 ovarian cancer cells after treatment with the 3D-NAF-1^44-67-6K^ peptide. **(a)** Representative confocal fluorescence images of mitochondria in SKOV-3 cells treated or untreated with the 3D-NAF-1^44-67-6K^ peptide. **(b)** Mitochondrial length was translated into skeleton binary images in the control (b. upper panel - without peptide) and with 50µM of the peptide treatment (b. lower panel). The mitochondrial length was measured in 25 cells from 8 field images and 156 mitochondria/cell in the control and 58 mito/cell in 50µM peptide treatment were measured using the Image J program. Mitochondrial length in the magnified images was measured using the free hand tool in image J program.

### 7. Studies to assess the toxicity of 3D-NAF-1^44-67-6K^ *in-vivo*

**Supplementary Table 2.**
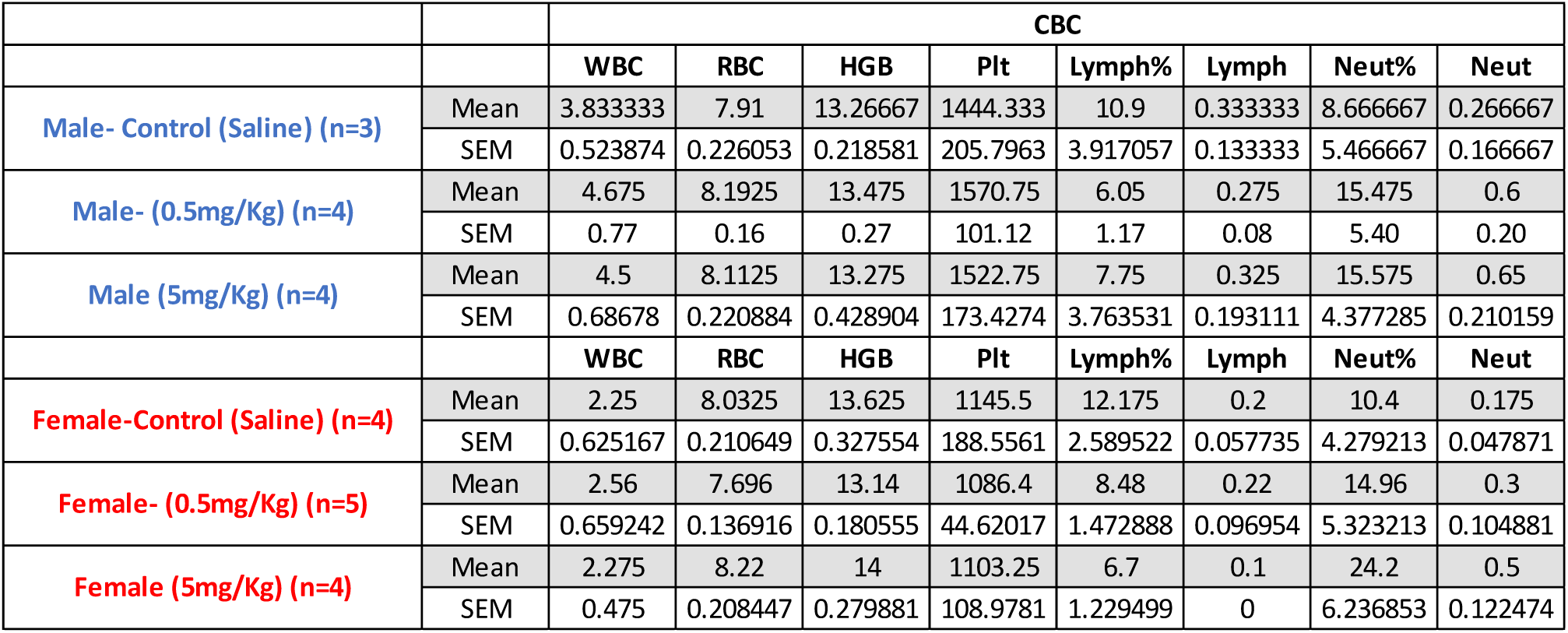
CBC results of male and female mice after treatment with/without the 3D-NAF-1^44-67-6K^ peptide. Treated mice were injected intraperitonially with 3D-NAF-1^44-67-6K^ using a concentration of 0.5mg/Kg or 5mg/Kg, while the control group was injected with Saline. Mice were injected 3 times a week for 4 weeks. Abbreviations; WBC: white blood cells, RBC: Red Blood cells, HGB: Hemoglobin, Plt: Platelets, Lymph: Lymphocytes cells, Neut: Neutrophils cells

**Supplementary Figure 8.**
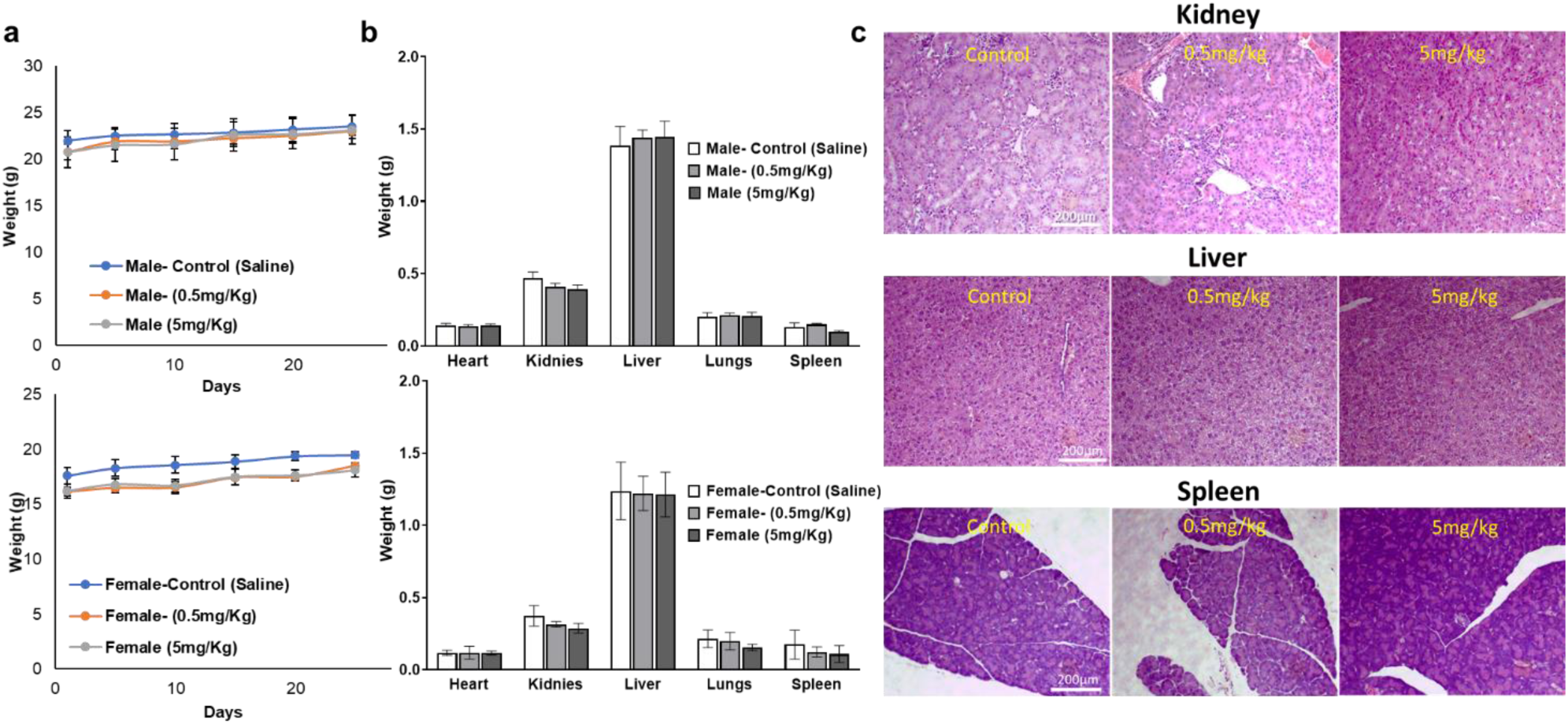
A toxicity study of 3D-NAF-1^44-67-6K^. **(a)** Male (upper panel) and female (lower panel) mice weight through the toxicity experiment period of the concentration of 0.5mg/Kg (orange), 5mg/Kg (grey) and saline control (blue). **(b)** Bar graphs showing the weight of the different organs harvested from Male (upper panel) and Female (lower panel) mice, treated or untreaded with the peptide at two different concentrations. **c)** representative H&E-stained sections of kidney, liver, and spleen, taken from the mice organs shown in (b), showing no differences between the tissues treated or untreated. Mice injected intraperitonially with 3D-NAF-1^44-67-6K^ using concentrations of 0.5mg/Kg or 5mg/Kg, while the control group was injected with Saline. Mice were injected 3 times a week for 4 weeks.

#### Supplementary methods for Simulation Methodology

Here we describe the setup for the simulations of the interactions of NAF-1^44-67-6K^ with the membranes (the simulations for NAF-1^44-67^ were run previously using a similar setup^4^). An initial structure of NAF-1^44-67-6K^ was built with AlphaFold^5, 6^. This predicted conformation was solvated in a water box, and a regular molecular dynamics trajectory was run for 200 ns. The final structure of the peptide in this aqueous system was used in the membrane simulations. Models for the normal and cancer membrane were built using 5 different lipids (Supplementary Table 1).

We built the two membranes with the Membrane Builder tool of CHARMM-GUI package^7, 8, 9^. During the building process we placed the peptide in the aqueous layer outside the membranes. The membranes were built with a total of 200 lipids. Enough water was added in the simulation box to fully solvate the membranes. Potassium and chloride ions were also added to neutralize the overall charge of the system and to obtain an ionic strength of 15 mM. The molecular dynamics simulations were performed with the program GROMACS^10^. We used the all-atom CHARMM36 force field model for the peptide and ions^11^, and the TIP3P model to describe the water molecules^12^. Periodic boundary conditions were applied in all directions. Electrostatics interactions were computed using the particle mesh Ewald method^13^, with a real space cutoff of 1.2 nm and a grid dimension of 0.12 nm. A 6-12 Lennard Jones potential was used to compute the van der Waals interactions with a cutoff of 1.2 nm with an additional switching function to make the force to decay to zero from 1.0 to 1.2 nm.

After running the standard CHARMM-GUI equilibration steps for 1.1 ns, we performed a 50-ns run at a constant pressure and temperature of 1 bar and 323.15 K, respectively. To control the temperature, we used v-rescaling with a stochastic term^14^, and for the pressure we used a Parrinello-Rahman coupling^15^, using a semi-isotropic scheme (the x and y directions fluctuated isotropically but the z-direction was allowed to fluctuate independently). In this trajectory, we restrained the distance of the center of mass of the peptide from the membrane surface to allow an unperturbed mixing of the phospholipids. After finishing that trajectory, we removed the restraint (making the peptide completely free to move) and run a 1.5 μs production trajectory with conformations saved every 100-ps for analysis. We used a time-step of 2 fs, constraining the water bonds with the SETTLE algorithm ^16^ and the hydrogen bonds for the rest of the system with the LINCS algorithm^17^.

## Description of Additional Supplementary Files

**Supplementary movies 1-3: 3D-NAF-1^44-67-6K^ dramatically decrease migration of SKOV-3 cells.** Time-lapse high-definition phase contrast movies of SKOV-3 cells migration treated with different concentrations of 3D-NAF-1^44-67-6K^: untreated control (Supplementary movie 1), 10µM 3D-NAF-1^44-67-6K^ (Supplementary movie 2) and 20µM 3D-NAF-1^44-67-6K^ (Supplementary movie 3). Cell phase contrast images were recorded every three hours for 96 hours.

**Supplementary movie 1:** SKOV-3 cells migration of untreated cells (control).

**Supplementary movie 2:** SKOV-3 cells migration of cells treated with 10µM 3D-NAF-1^44-67-6K^.

**Supplementary movie 3:** SKOV-3 cells migration of cells treated with 20µM 3D-NAF-1^44-67-6K^.

